# AOPGraphExplorer 2.0: An Interactive Graph-Based Platform for Multi-Domain Mechanistic Annotation and Exploration of Adverse Outcome Pathways

**DOI:** 10.64898/2026.02.27.708648

**Authors:** Asmaa A. Abdelwahab, Barry Hardy

## Abstract

The Adverse Outcome Pathway (AOP) framework is a cornerstone of modern mechanistic toxicology, providing a structured representation of causal biological events linking molecular initiating events to adverse health outcomes. However, the practical exploration and interpretation of AOPs remain challenging due to the fragmentation of mechanistic knowledge across heterogeneous biological databases and the limited availability of integrated, interactive tools.

Here, we present AOPGraphExplorer 2.0, an interactive graph-based platform for the visualization, annotation, and analysis of AOP networks derived from AOP-Wiki. This new version introduces a scalable, modular architecture that integrates multi-domain mechanistic annotations, including biological processes, molecular entities, anatomical context, diseases, and stressors, directly into AOP graphs.

AOPGraphExplorer 2.0 enables dynamic filtering of AOP networks based on weight-of-evidence and quantitative understanding scores, supports AOP-, Key Event-, and keyword-centric queries, and generates exportable interactive HTML and machine-readable JSON representations for documentation, sharing, and computational reuse. In addition, the platform provides automated network statistics and annotation coverage summaries, supporting transparent and reproducible analysis.

By bridging AOP-Wiki with external biomedical knowledge resources in a unified graph framework, AOPGraphExplorer 2.0 transforms AOP exploration into a multi-domain systems-level analysis workflow. The platform supports hypothesis generation, mechanistic interpretation, and evidence-based decision-making in toxicology, pharmacology, and risk assessment.

## 1. Introduction

### 1.1 The Adverse Outcome Pathway (AOP) framework

The Adverse Outcome Pathway (AOP) framework is an analytical construct for organizing mechanistic toxicology knowledge into a biologically plausible, evidence-supported chain of events that links an initial molecular perturbation to an adverse outcome relevant to risk assessment and regulatory decision-making (Ankley *et al*., 2010; Villeneuve *et al*., 2014; OECD, 2018). An AOP is typically anchored by a Molecular Initiating Event (MIE) (the first interaction/perturbation at the molecular level), followed by one or more measurable Key Events (KEs) across levels of biological organization, and culminates in an Adverse Outcome (AO) at an apical level of biological relevance. The causal and predictive linkages between upstream and downstream KEs are captured as Key Event Relationships (KERs), which provide the scientific basis for inference and extrapolation from early events to later outcomes (OECD, 2018).

A central principle of the framework is modularity: KEs and KERs are reusable building blocks that can be shared across multiple AOPs, enabling the assembly of AOP networks to reflect real-world biological complexity and support cross-pathway reasoning (Villeneuve *et al*., 2014; Knapen *et al*., 2018). To promote transparency, consistency, and community-driven curation, the OECD AOP Knowledge Base—particularly AOP-Wiki—provides a standardized narrative structure and supporting metadata, and has increasingly been made more computable through ontology-based annotations and semantic representations that facilitate integration with external biological and chemical resources (OECD, 2018; Ives *et al*., 2017; Martens *et al*., 2022).

Because AOPs connect mechanistic evidence to outcomes of regulatory concern, they are widely used to support New Approach Methodologies (NAMs) and Integrated Approaches to Testing and Assessment (IATA) by providing a coherent structure for integrating *in vitro*, in silico, and emerging data streams (e.g., high-throughput and omics evidence) into fit-for-purpose decision contexts (Tollefsen *et al*., 2014; Kleinstreuer *et al*., 2016; Perkins *et al*., 2015).

### 1.2 Challenges in AOP data exploration

AOP-Wiki serves as the primary repository for curated AOP knowledge; however, several challenges limit its effective use for mechanistic reasoning: AOP-Wiki, the main module of the OECD AOP Knowledge Base, serves as the primary community repository for curated AOP descriptions and supporting evidence (Villeneuve *et al*., 2014; Martens*et al*., 2022; AOP-Wiki, n.d.). While its page-centric structure is effective for human curation, systematic reuse for mechanistic reasoning and computational analysis remains challenging, especially when AOPs must be interpreted as interconnected networks and linked to external biomedical and toxicological resources (Knapen *et al*., 2018; Martens *et al*., 2022; Mortensen *et al*., 2025).

Key challenges include:

- **Fragmented biological context across resources**. Interpreting AOPs often requires linking Key Events (KEs) to genes/proteins, pathways, phenotypes, tissues, and chemicals distributed across multiple databases and ontologies, which is not consistently available as machine-readable context in AOP-Wiki entries (Ives *et al*., 2017; Martens *et al*., 2018; Mortensen *et al*., 2022).
- **Incomplete structuring of mechanistic information**. Important mechanistic details may remain embedded in free text rather than standardized fields, limiting reliable downstream querying and integration (Martens *et al*., 2022; Virvilis & Lekka, 2025).
- **Limited network-level exploration in the native interface**. Although AOPs can be read individually, exploring multi-AOP causal chains, alternative routes, and emergent pathways is difficult without network-centric views and workflow support (Knapen *et al*., 2018; Yarar *et al*., 2025).
- **Non-trivial extraction and integration overhead**. Automatic parsing and systematic reuse of AOP-Wiki content is well-recognized as challenging, motivating parallel efforts such as semantic representations (AOP-Wiki RDF) and integrated resources (e.g., AOP-DB/AOP-DB RDF) (Martens *et al*., 2022; Pittman *et al*., 2018; Mortensen *et al*., 2022).
- **High barrier for non-technical users**. Advanced reuse frequently requires RDF/SPARQL, identifier harmonization, or specialized analysis environments, which increases effort for routine exploration and hypothesis generation (Martens *et al*., 2022; Mortensen *et al*., 2025; Yarar *et al*., 2025).

A growing ecosystem of community tools extends AOP-Wiki exploration and visualization, including interactive network viewers and knowledge-driven navigation (e.g., Biovista Vizit, Wiki Kaptis, AOP Mapper, AOP-networkFinder), literature-mining support (AOP-helpFinder), semantic exports (AOP-Wiki RDF), and omics-oriented analytics (BMDx, AOPfingerprintR) (AOP-Wiki, n.d.; Virvilis & Lekka, 2025; Kane *et al*., 2023; Jornod *et al*., 2022; Martens *et al*., 2022; Mortensen *et al*., 2021; Serra *et al*., 2020; Yarar *et al*., 2025; Serra & Fratello, 2025). However, these tools are typically accessed as separate services with heterogeneous input/output formats and varying technical prerequisites (e.g., configuration, specialized runtimes, RDF/SPARQL expertise, or post-processing for cross-database linking), making it non-trivial to benefit from them in a unified, low-effort workflow (Martens *et al*., 2022; Mortensen *et al*., 2025; Yarar *et al*., 2025).

To address this gap, AOPGraphExplorer 2.0 aims to provide an easy-to-use web framework that lowers the barrier for AOP data exploration by offering interactive visualization on top of AOP-Wiki content, multi-AOP/network-scale navigation, and streamlined access to AOP–biomedical–omics linkouts and AI/ML-ready graph representations, with minimal setup, configuration, or specialist knowledge required.

### 1.3 Motivation and prior work

AOPGraphExplorer 1.0 introduced interactive, graph-based visualization of AOP structures, enabling users to navigate MIE–KE–AO relationships and explore connected KEs and KERs beyond static AOP-Wiki pages (Ali & Hardy, 2025). However, the first version provided limited support for systematic, multi-domain annotations (e.g., pathways, genes/proteins, tissues, disease links, and omics-relevant context), offered restricted evidence-aware filtering, and did not provide reusable, exportable graph representations designed for downstream analysis and integration into external workflows.

As AOP-centric research grows in scale and moves toward increasingly data-driven use cases (e.g., IATA/NAM-enabled decision contexts, quantitative AOP development, and network-level reasoning), there is an increasing need for a unified platform that (i) integrates curated AOP knowledge with interoperable mechanistic annotations, (ii) supports transparent, evidence-aware prioritization using established AOP descriptors (including weight-of-evidence and quantitative understanding where available), and (iii) enables reproducible and shareable network analysis through machine-actionable exports aligned with FAIR principles.

### 1.4 AOPGraphExplorer 2.0 Contributions

As summarized in the graphical abstract (Fig. 1), AOPGraphExplorer 2.0 delivers the following advances for exploring and analyzing AOP knowledge:

**Figure 1.**
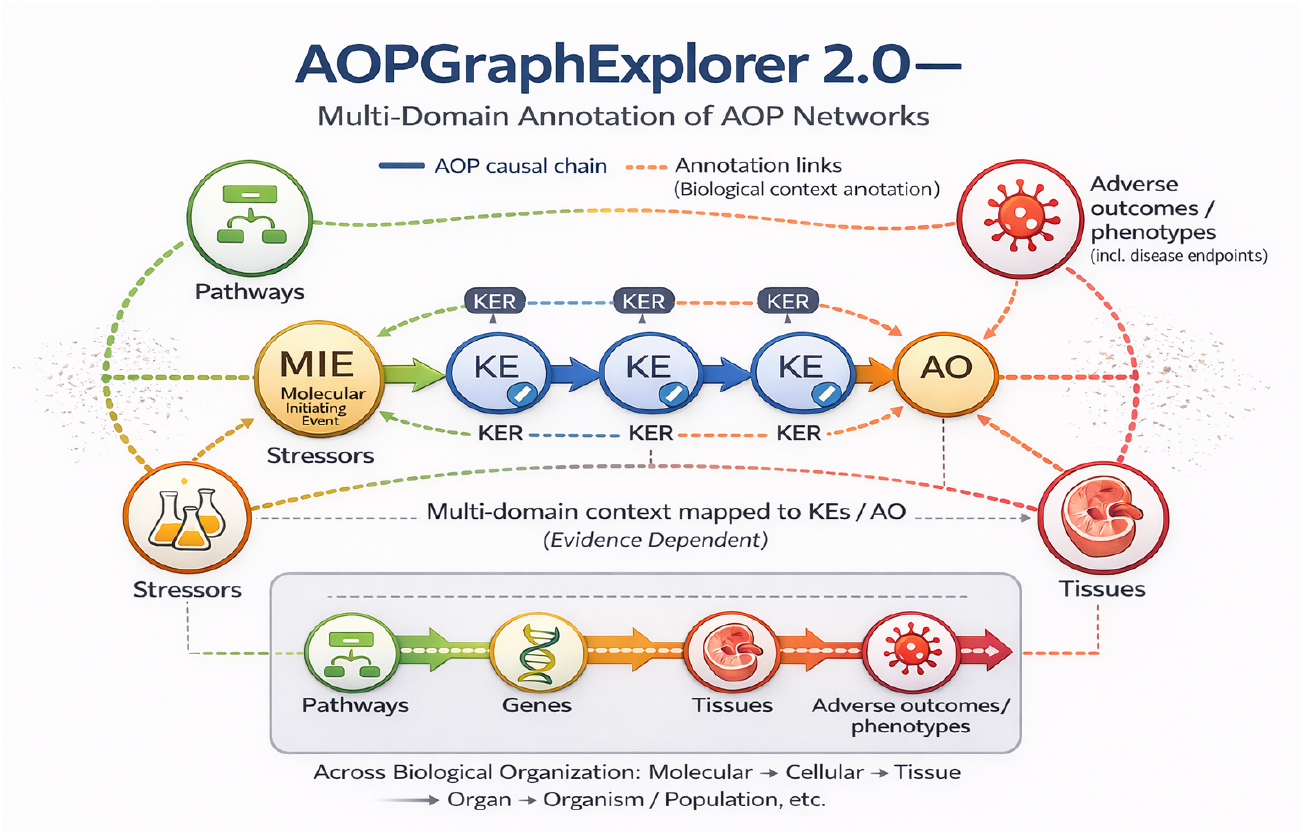
Graphical abstract of AOPGraphExplorer 2.0. AOPGraphExplorer 2.0 represents AOP knowledge as a directed network in which the core AOP causal chain (MIE → KE(s) → AO) is connected by Key Event Relationships (KERs). The platform extends this core network with evidence-dependent multi-domain annotation links that map biological and contextual entities (e.g., pathways/processes, genes/proteins, chemicals/stressors, tissues/anatomy, and adverse outcomes/phenotypes including disease endpoints) onto KEs and AOs. Solid edges denote causal AOP relationships, while dashed edges denote contextual annotation links.

- **Multi-domain mechanistic annotation of Key Events**, linking KEs to biological and contextual entities (e.g., pathways, proteins/genes, chemicals, tissues, diseases, and population/context descriptors).
- **A unified graph schema** that integrates canonical AOP elements (MIE–KE–AO and KERs) with modular enrichment layers to support scalable extension and reuse.
- **Evidence-aware filtering and prioritization**, enabling users to rank and subset networks using **weight-of-evidence** and **quantitative understanding** descriptors (as captured in AOP-Wiki where reported).
- **Interactive graph visualization with enrichment-layer controls**, allowing users to toggle annotation layers and tailor views to specific analysis questions.
- **Reusable export formats**, including self-contained **HTML** for sharing and **JSON** for downstream computational workflows.
- **Automated network summaries**, providing basic network statistics and annotation coverage reports to support transparency, traceability, and reproducibility.
- **Flexible search and entry points**, supporting keyword-based exploration as well as AOP-centric and KE-centric workflows.

## 2. Data Sources and System Architecture

### 2.1 AOP-Wiki data ingestion

AOPGraphExplorer 2.0 ingests structured content from AOP-Wiki using a hybrid strategy that combines (i) periodic cached snapshots of the official XML releases and (ii) tabular exports (TSV) for high-use entities, including Key Events (KEs), Key Event Relationships (KERs), and Key Event Components. This approach balances completeness (XML) with efficient downstream processing (TSV).

During ingestion, the pipeline parses, validates, and normalizes core AOP entities and metadata, including: AOP identifiers and titles; Key Events and their roles (MIE, intermediate KE, AO); Key Event Relationships with directionality and confidence descriptors (weight-of-evidence and quantitative understanding, where reported); stressors and chemical descriptors; and biological applicability fields such as taxonomy, sex, and life stage. All inputs are cached as versioned quarterly snapshots, enabling reproducible analyses by ensuring that interactive sessions and exports can be traced to a stable data release.

### 2.2 Multi-domain annotation sources

To support mechanistic interpretation beyond linear AOP sequences, AOPGraphExplorer 2.0 enriches AOP networks with multi-domain biological and contextual entities linked to Key Events (KEs) and, where applicable, Adverse Outcomes (AOs). Annotation nodes and links are derived primarily from structured fields in AOP-Wiki XML exports and are complemented with standardized cross-references to external knowledge resources (Table 1). The integrated domains include biological processes and pathways (e.g., GO, KEGG, Reactome, WikiPathways), genes/proteins and molecular entities (e.g., UniProt, Ensembl, Protein Ontology), chemical and stressor identifiers (e.g., ChEBI, PubChem, DSSTox), anatomy/tissue context (e.g., Uberon, FMA, BRENDA Tissue Ontology), and disease/phenotype descriptors (e.g., MeSH, CTD, HPO/MP, PCO). Annotation is implemented using a consistent mapping strategy based on ontology identifiers and normalized cross-references. Provenance is preserved throughout via source identifiers and outbound database links, enabling traceable inspection of the evidence underlying each annotation.

**Table 1.** Multi-domain annotation sources used by AOPGraphExplorer 2.0. The table summarizes external resources used to standardize and enrich KE/AO context via identifiers and controlled vocabularies.

| Annotation domain | Resource | What it contributes | Citation |
| --- | --- | --- | --- |
| **Processes & pathways** | Gene Ontology (GO) | Standardized biological process/function terms for KE context and aggregation | (The Gene Ontology Consortium, 2026) |
| **Processes & pathways** | KEGG | Curated pathway identifiers and pathway context | (Kanehisa & Goto, 2000) |
| **Processes & pathways** | Reactome | Curated human pathway knowledgebase and pathway cross-references | (Milacic *et al.*, 2024) |
| **Processes & pathways** | WikiPathways | Community-curated pathways and pathway identifiers | (Agrawal *et al.*, 2024) |
| **Genes/proteins** | UniProt | Protein identifiers and cross-references for molecular entity mapping | (The UniProt Consortium, 2025) |
| **Genes/proteins** | Ensembl | Gene-centric identifiers and genome annotation cross-references | (Dyer *et al.*, 2025) |
| **Genes/proteins** | Protein Ontology (PRO) | Ontology for protein forms/complexes to standardize protein entities | (Natale *et al.*, 2017) |
| **Chemicals/stressors** | ChEBI | Standard chemical entity identifiers and ontology terms | (Malik *et al.*, 2026) |
| **Chemicals/stressors** | PubChem | Chemical identifiers and compound metadata for stressor/chemical mapping | (Kim *et al.*, 2025) |
| **Chemicals/stressors** | DSSTox | Curated chemical identifiers supporting computational toxicology | (Grulke *et al.*, 2019) |
| **Anatomy/tissue** | Uberon | Cross-species anatomy ontology for tissue/anatomy context | (Mungall *et al.*, 2012) |
| **Anatomy/tissue** | Foundational Model of Anatomy (FMA) | Human anatomy reference ontology for anatomical mapping | (Rosse & Mejino, 2003) |
| **Anatomy/tissue** | BRENDA Tissue Ontology (BTO) | Tissue terms for enzyme/tissue source context and mapping | (Gremse *et al.*, 2011) |
| **Disease/phenotype** | MeSH | Controlled vocabulary for biomedical topics and disease-related descriptors | (National Library of Medicine, n.d.) |
| **Disease/phenotype** | CTD | Curated chemical–gene–disease associations to support disease context links | (Davis *et al.*, 2025) |
| **Disease/phenotype** | Human Phenotype Ontology (HPO) | Human phenotype descriptors for phenotype-level mapping | (Gargano *et al.*, 2024) |
| **Disease/phenotype** | Mammalian Phenotype Ontology (MP) | Mammalian phenotype terms to support cross-species phenotype mapping | (Bello *et al.*, 2025) |
| **Population/context** | Population and Community Ontology (PCO) | Population-level descriptors for context/population annotations | (Population and Community Ontology Developers, n.d.) |

### 2.3 System architecture

AOPGraphExplorer 2.0 follows a modular architecture designed to support reproducible graph assembly, evidence-aware exploration, and interactive visualization (Fig. 2). The system is organized into sequential components: (1) data loading and snapshot caching, (2) identifier normalization and schema harmonization, (3) construction of the core AOP causal graph (MIE–KE–AO connected by KERs), (4) optional enrichment with multi-domain annotation layers, (5) evidence-based filtering and user-driven subsetting, and (6) interactive visualization and export for downstream reuse. This separation of concerns enables independent updating of source snapshots and annotation layers while maintaining stable graph identifiers and consistent user-facing outputs.

**Figure 2.**
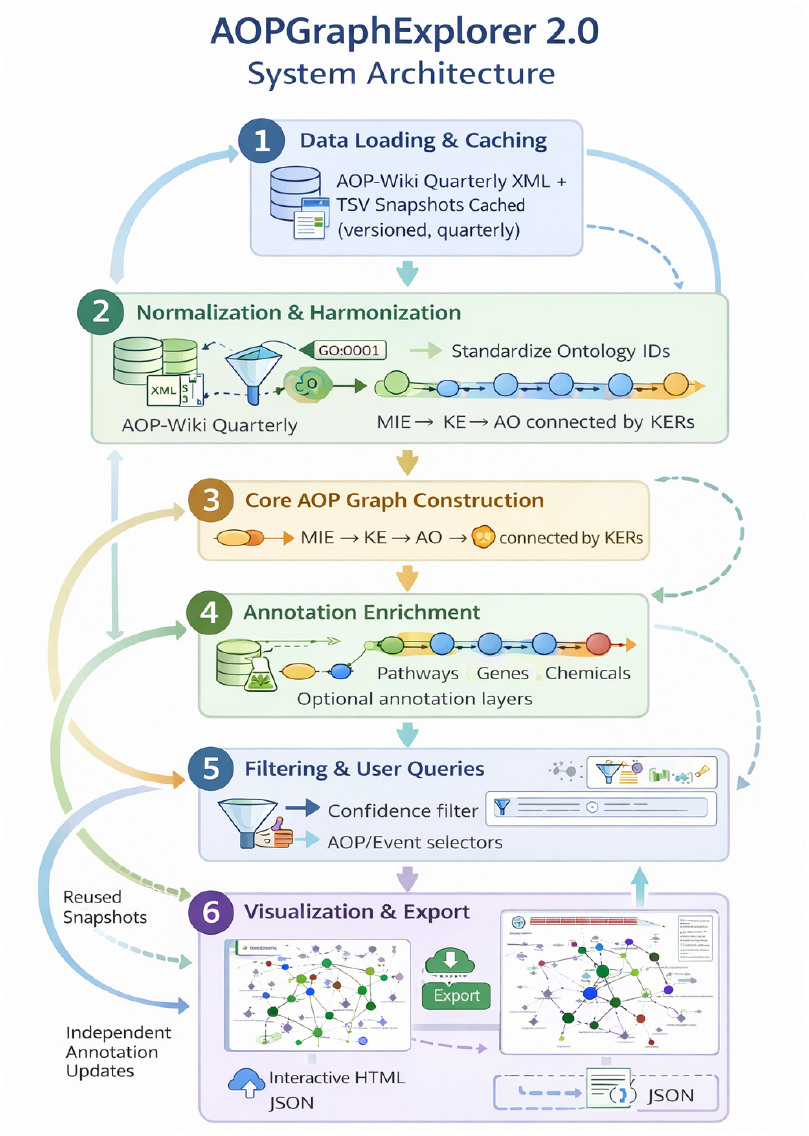
AOPGraphExplorer 2.0 system architecture. The platform is implemented as a modular pipeline comprising (1) AOP-Wiki data loading and quarterly snapshot caching, (2) identifier normalization and schema harmonization, (3) core AOP graph construction (MIE–KE–AO linked by KERs), (4) optional multi-domain annotation enrichment, (5) evidence-based filtering and user-driven querying, and (6) interactive visualization and export (e.g., screenshots/HTML views and machine-readable JSON). The design supports reproducibility through versioned snapshots and enables independent updates of annotation layers without changing stable graph identifiers.

## 3. Methods and Implementation

### 3.1 Graph construction

AOPGraphExplorer 2.0 represents AOP-Wiki content as a directed, typed graph that separates the **core AOP causal structure** from **contextual annotation layers**. The core network comprises nodes corresponding to AOP elements, including Molecular Initiating Events (MIEs), intermediate Key Events (KEs), and Adverse Outcomes (AOs), as well as AOP entities themselves and stressors where available. To enable multi-domain interpretation, the graph may be extended with additional nodes representing annotated biological entities, such as biological processes/pathways, proteins/genes, chemicals, anatomy/tissues, and disease/phenotype terms. Directed edges encode Key Event Relationships (KERs) and thus capture the causal backbone of the AOP structure, while additional edge types encode AOP membership, stressor–event associations, and evidence-linked annotation relations between events and contextual entities. Node and edge identifiers are normalized and stabilized to ensure consistent referencing across user sessions, exports, and downstream analyses.

### 3.2 Annotation integration pipeline

For each KE (and AO where applicable), AOPGraphExplorer 2.0 integrates structured biological context derived from AOP-Wiki exports into the graph as evidence-linked annotation layers. The pipeline parses relevant XML fields describing biological context and supporting metadata, then maps biological processes and actions to standardized ontology terms (e.g., GO/NBO), enabling consistent search, aggregation, and reuse across AOPs. Molecular entities (proteins/genes), chemicals, anatomy/tissue terms, and disease/phenotype descriptors are associated using external identifiers where available, and are incorporated as separate annotation nodes connected to KEs/AOs via typed semantic edges. Provenance is preserved throughout, including source identifiers and outbound database links to support validation and traceability. Importantly, annotation links are treated as contextual associations rather than causal assertions and can be enabled or disabled dynamically without modifying the underlying core AOP causal network.

### 3.3 Evidence-based filtering

To support evidence-aware exploration, AOPGraphExplorer 2.0 provides filtering mechanisms that operate on both the structure of the network and confidence assessments associated with KERs. Users can restrict displayed networks based on Weight-of-Evidence (WoE) and Quantitative Understanding scores, as well as subset the network by selected AOPs, events (MIE/KE/AO), or stressors. The platform also supports restricting views to complete versus incomplete causal sequences, depending on data availability. Filtering is applied prior to visualization and summary generation so that both the displayed graph and computed statistics reflect the selected evidence criteria.

### 3.4 Visualization engine

AOPGraphExplorer 2.0 provides an interactive visualization layer for exploratory analysis of multi-AOP networks. The interface supports physics-enabled layouts that help users interpret complex graph topologies, and it applies consistent visual encodings to distinguish AOP event categories (MIE, KE, AO) and integrated annotation entity types (e.g., process, protein, chemical, anatomy, disease, phenotype). Core causal edges (KERs) are visually separated from enrichment/annotation links using distinct styling (e.g., solid vs. dashed lines), improving interpretability and reducing the risk of conflating contextual associations with causal relationships.

To highlight cross-AOP convergence, edge thickness encodes the number of shared occurrences of that connection across pathways (i.e., how many AOPs contain the same source–target relationship). The edge tooltip/properties list the corresponding AOP ID(s) supporting that shared connection, enabling immediate traceability back to the underlying pathways. Similarly, node size and node border thickness scale with the number of pathways in which the node is shared, emphasizing hubs and commonly reused events or annotations. The exact count of AOPs sharing each node is also included in the node tooltip/properties, making the degree of reuse explicit.

Interactive, HTML-based tooltips expose node- and edge-level metadata and provide clickable links to relevant external resources (e.g., AOP-Wiki for events and relationships, Reactome/AmiGO for processes, UniProt/HGNC for proteins, and ChEBI for chemicals), supporting transparent navigation of evidence and context. Legends update dynamically when annotation layers are toggled, and graph views can be exported (e.g., screenshots) to support reporting and reuse.

### 3.5 Network statistics and summaries

To facilitate reporting and comparative analysis, AOPGraphExplorer 2.0 automatically computes descriptive statistics for the active network state. These include node and edge counts by type, annotation coverage metrics (e.g., the proportion of KEs connected to process/protein/anatomy/disease terms), and distributions of KER confidence assessments (WoE and quantitative understanding). The platform additionally computes basic connectivity descriptors such as connected components and overall graph size, and provides summaries of stressor participation and recurring annotated biological themes across selected AOPs. All computed statistics are exportable for integration into manuscripts, technical reports, and supplementary materials.

### 3.6 Parkinson’s case study workflow

Guided by the AOPGraphExplorer 2.0 concept of integrating the core AOP causal chain with evidence-dependent multi-domain annotations (Graphical Abstract; Fig. 1), we conducted a Parkinson’s disease (PD)–motivated case study to demonstrate mechanistic exploration of AOP networks. Candidate AOPs were retrieved from AOP-Wiki using keyword queries (e.g., “parkinson”, “neurodegeneration”, “dopaminergic”, “mitochondrial”, “neuroinflammation”) and were curated by screening titles and abstracts to retain neurotoxicity/neurodegeneration-relevant pathways. A combined network was then assembled using Molecular Initiating Events (MIEs), Key Events (KEs), Adverse Outcomes (AOs), and Key Event Relationships (KERs). Mechanistic annotation layers (process/pathway, protein, chemical, anatomy, disease/phenotype) were enabled to map contextual evidence onto KEs/AOs, consistent with the multi-domain annotation model (Fig. GA). To assess robustness and support evidence-aware interpretation, networks were examined under confidence filtering of KERs (weight-of-evidence and quantitative understanding scores), comparing the full annotated network to a higher-confidence subnetwork (workflow illustrated in Fig. 3A–C).

**Figure 3.**
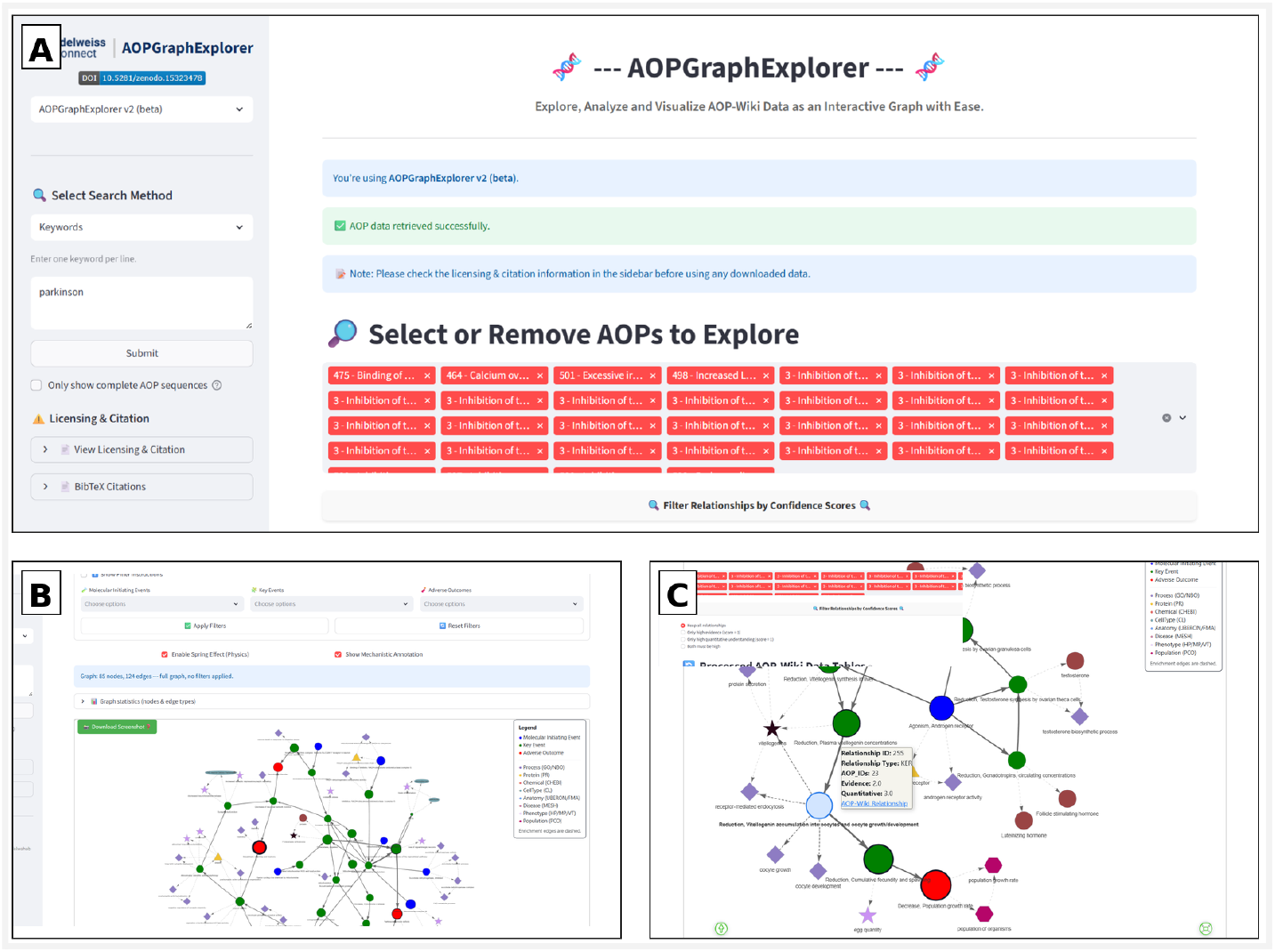
Parkinson’s case study workflow in AOPGraphExplorer 2.0. (A) Keyword-based retrieval and selection of candidate AOPs from AOP-Wiki using the PD query (illustrated with “parkinson”); (B) Full network visualization with mechanistic annotations enabled, showing the combined AOP graph and the annotation legend (core AOP elements and enrichment entity types); (C) Evidence-aware exploration through confidence-based filtering and inspection of a Key Event Relationship (KER) tooltip reporting evidence and quantitative understanding scores, illustrating how users can prioritize higher-confidence subnetworks.

## 4. Results

### 4.1 Integrated AOP graph overview

Using the current AOP-Wiki snapshot, AOPGraphExplorer 2.0 constructs an integrated, directed network that combines canonical AOP structure (MIE–KE–AO linked by KERs) with optional multi-domain annotation layers. Across the retrieved content, the resulting graphs include large numbers of KEs and KERs, and expose evidence descriptors (e.g., weight-of-evidence and quantitative understanding, where reported) alongside network topology. When annotation layers are enabled, KEs and AOs are linked to contextual entities spanning biological processes/pathways, proteins/genes, chemicals/stressors, anatomy/tissues, and disease/phenotype descriptors. In addition to interactive exploration, the platform provides summary statistics describing node/edge composition, annotation coverage, and the distribution of evidence-related descriptors, supporting transparent reporting and comparison across query results.

### 4.2 Use case demonstration - Parkinson’s-focused AOP exploration

To demonstrate practical use in a neurotoxicity context, we conducted a Parkinson’s disease (PD)–motivated case study based on keyword-driven retrieval and curation of PD-relevant AOPs (workflow in Fig. 2; conceptual mapping in Fig. 1/Graphical Abstract). The curated AOP set was assembled into an integrated, directed network that preserves the MIE–KE–AO structure and KER directionality, and was subsequently enriched with optional mechanistic annotation layers.

In the exported PD network, the causal core comprised 12 molecular initiating events (MIEs), 24 key events (KEs), and 3 adverse outcomes (AOs), connected by KER edges. Causal relationships remain traceable at the edge level via interactive tooltips that report the AOP-Wiki relationship identifier, AOP_ID membership, and, when available, weight-of-evidence and quantitative understanding scores. This design enables rapid comparison between the full network and higher-confidence mechanistic routes by supporting filtering and inspection without loss of provenance.

A principal advantage of the integrated view is that multiple upstream perturbations converge on PD-relevant downstream outcomes. In this network, convergence is mediated by a small set of highly connected KEs, including mitochondrial dysfunction, intracellular calcium overload, oxidative stress, and proteostasis disruption, ultimately linking to degeneration of dopaminergic neurons in the nigrostriatal pathway and Parkinsonian motor deficits (both represented explicitly as downstream PD-relevant endpoints). Shared-node metadata further highlights cross-AOP reuse of central events (e.g., dopaminergic neuron degeneration and motor deficits shared across multiple AOPs), supporting prioritization of common mechanistic bottlenecks for follow-up assessment.

Mechanistic interpretability is further strengthened by the multi-domain enrichment layer (46 annotation nodes in this example), which links events to biological processes (e.g., glutamatergic/NMDA receptor activity, calcium ion transport, mitochondrial components and electron transport), phenotypes (e.g., oxidative stress, loss of dopaminergic neurons, dementia-related phenotypes), disease descriptors (e.g., Parkinsonian disorders), and relevant cell types (e.g., CNS neurons, astrocytes, microglia). This contextual overlay provides biologically meaningful entry points for exploration—for example, selecting an event to highlight its local neighborhood and inspecting linked metadata via tooltips—thereby supporting domain-driven interpretation of causal routes.

As shown in Figure 4, the integrated PD network reveals several representative causal routes that converge on PD-like endpoints. These include: (i) mitochondrial inhibition → mitochondrial dysfunction → dopaminergic neurodegeneration → motor deficits, exemplified by complex I inhibitor binding leading to complex I inhibition and increased mitochondrial dysfunction before converging on dopaminergic neuron degeneration and Parkinsonian motor deficits; (ii) excitotoxicity → calcium overload → calpain/caspase activation → cell death → motor deficits, reflecting a high-confidence glutamatergic/NMDA-driven route; and (iii) iron accumulation/oxidative stress → misfolded proteins/proteostasis stress → apoptosis → cell death → motor deficits, linking oxidative and proteostasis mechanisms to PD-like adverse outcomes. Across the curated PD AOP set, the network highlights two dominant convergence motifs toward PD-like endpoints: (1) mitochondrial bioenergetic disruption/ROS and (2) glutamatergic excitotoxicity/calcium overload, with downstream coupling to proteostasis stress and apoptotic cell death prior to dopaminergic neuron degeneration and motor impairment.

**Figure 4.**
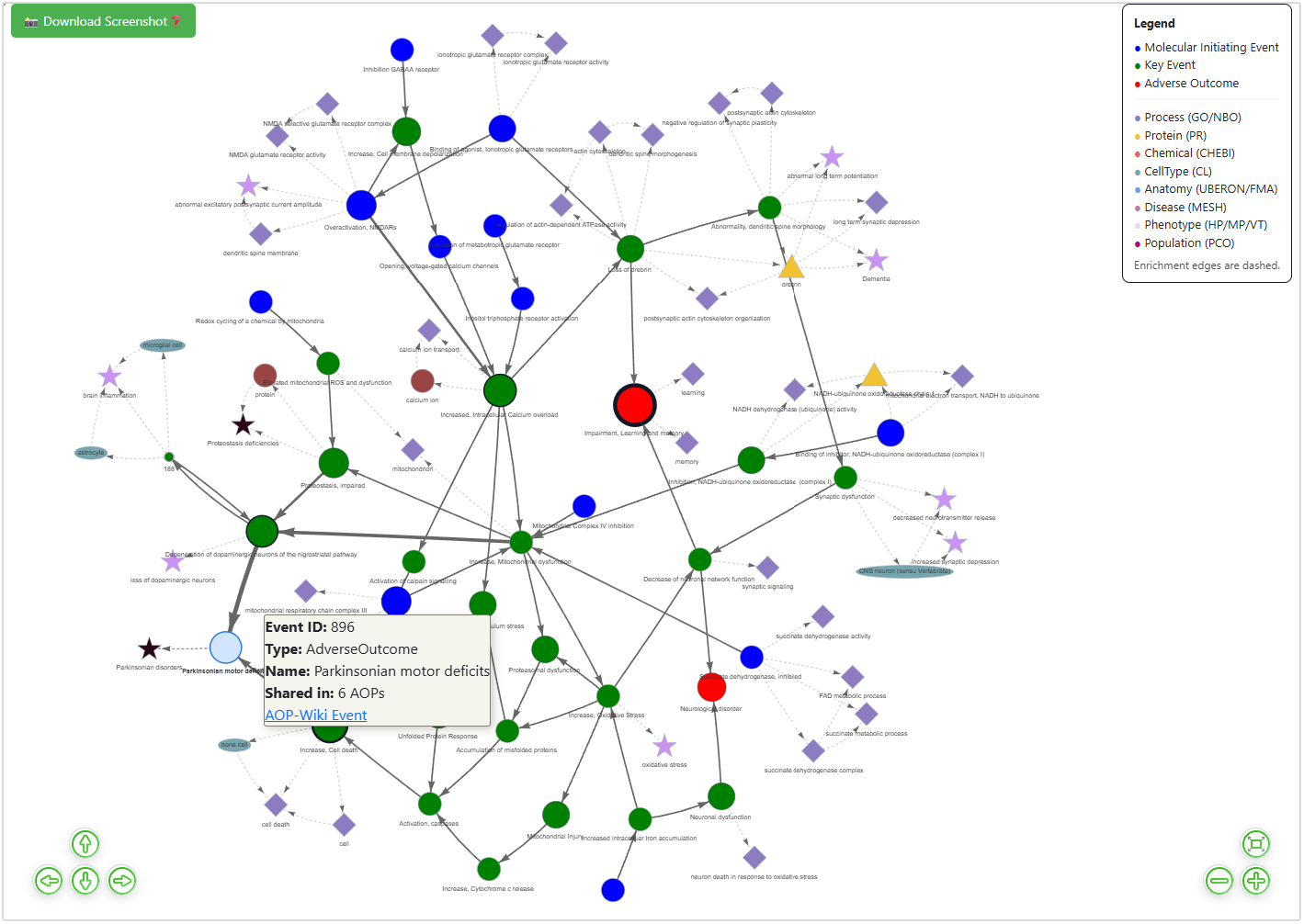
Parkinson’s disease (PD)–motivated integrated AOP network generated with AOPGraphExplorer 2.0. The directed causal core (solid arrows) connects molecular initiating events (MIEs; blue), key events (KEs; green), and adverse outcomes (AOs; red) through key event relationships (KERs), while dashed edges represent optional enrichment links to mechanistic context (processes, proteins, chemicals, phenotypes, diseases, anatomy, cell types, and populations; node symbols/colors as indicated in the legend). Node size and border thickness reflect cross-AOP reuse of events, and causal edge width reflects the number of AOPs supporting a given KER. Interactive tooltips expose event/relationship identifiers, AOP membership (AOP_IDs), and available confidence metrics, with direct links to the corresponding source entries (e.g., AOP-Wiki).

Because the initial AOP set was assembled via keyword-based retrieval, the composition and specificity of the resulting PD network are sensitive to the choice of search terms. Iterative refinement of keywords, guided by the mechanistic scope of the case study and by inspection of retrieved AOP titles, KEs, and stressors, can improve precision by reducing off-target pathways and improve recall by capturing relevant synonyms and related descriptors. Accordingly, keyword selection should be treated as a tunable step that can be tailored to the scientific focus and intended use of the case study, enabling more targeted network construction and downstream interpretation.

Overall, this case study illustrates how multi-domain enrichment can improve interpretability and prioritization relative to viewing AOPs as linear sequences alone, while preserving traceability to source evidence through direct links and provenance metadata embedded in node- and edge-level tooltips. Use case data are available via Zenodo (DOI: 10.5281/zenodo.18732647), and step-by-step guidance is provided in the **Supplementary Materials**.

## 5. Discussion

### 5.1 Significance

AOPGraphExplorer 2.0 enables AOP knowledge to be explored as an integrated, evidence-aware network rather than as isolated linear pathways. By unifying AOP-Wiki causal structure with optional multi-domain context layers, the platform reduces manual effort required to connect AOP elements to relevant biological entities and external resources. This supports transparent mechanistic interpretation, hypothesis generation, and targeted prioritization of KEs/KERs for downstream assessment, while maintaining reproducibility through snapshot-based data handling and stable identifiers.

### 5.2 Limitations

Several limitations should be considered. First, annotation is constrained by the availability and structure of AOP-Wiki metadata and the completeness of cross-references; consequently, annotation density and quality can vary substantially between AOPs. Second, mappings to controlled vocabularies may remain incomplete when source terms are ambiguous, underspecified, or lack stable identifiers. Third, evidence-aware filtering and network prioritization reflect the confidence descriptors available in AOP-Wiki; missing or inconsistently reported scores can limit comparability across KERs. Finally, the platform does not aim to infer causality beyond curated KERs; annotation links are contextual and should not be interpreted as causal edges.

### 5.3 Future directions

Planned developments include (i) AI-assisted support for KE annotation, similarity analysis, and candidate cross-linking across AOPs, (ii) deeper integration with AOPOntology and semantic web standards to improve interoperability and machine-actionability, (iii) network-level similarity, clustering, and motif-based analyses across AOP collections, and (iv) coupling with quantitative toxicokinetic and dynamic models to support more quantitative, context-specific interpretation of pathways and outcomes.

## 6. Conclusion

AOPGraphExplorer 2.0 provides an extensible, interactive platform for evidence-aware, multi-domain exploration of AOP knowledge. By representing AOP-Wiki content as a unified graph and optionally enriching core AOP structure with contextual biological annotations, the platform improves transparency, supports mechanistic interpretation, and enables reproducible workflows for AOP exploration and prioritization in risk and safety assessment contexts.

## 7. Availability

- **AOPGraphExplorer releases (Zenodo):** https://zenodo.org/records/18728687
- **Online deployment:** https://aopgraphexplorer.edelweissconnect.com/
- **User guide:** A complete video tutorial is available via the online deployment page.

## 8. Author Contributions

**Asmaa A. Abdelwahab:** Conceptualization, system design, software development, data analysis, manuscript writing

**Barry Hardy:** Supervision, conceptual guidance, manuscript review and revision

## 9. Funding & Acknowledgements

### Funding

Not applicable.

## Acknowledgements

We thank **[Jeff Wiseman, Edelweiss Connect GmbH]** for applying AOPGraphExplorer 2.0 in his ongoing research and for providing valuable feedback on functionality and usability.

## 10. Supplementary materials

Supplementary information is available in the Supplementary Materials document: Supplementary Materials

