## Supplementary materials for "AOPGraphExplorer 2.0: An Interactive Graph-Based Platform for Multi-Domain Mechanistic Annotation and Exploration of Adverse Outcome Pathways"

### S1. Objective and scope

This case study demonstrates how **AOPGraphExplorer 2.0** supports interpretation of a **Parkinson's disease (PD)-motivated multi-AOP network** by combining a directed causal core (MIE–KE–AO) with optional mechanistic annotation layers (processes, phenotypes, cell types, diseases, proteins, and chemicals). The content below is derived directly from the **provided HTML export**.

### S2. Input data and curated AOP set (as represented in the export)

A PD-relevant AOP subset was assembled via keyword-based retrieval. In the exported network, causal KER edges reference the following **AOP-Wiki AOP identifiers**:
**AOP IDs:** 3, 464, 475, 498, 501, 587, 588, 589, 593.

AOP 464 dominates the causal backbone in this export (appearing in the largest number of KER edge metadata entries), indicating that one AOP (or a closely related group of KERs) provides a substantial portion of the mechanistic chain(s) connecting upstream perturbations to PD-like outcomes.

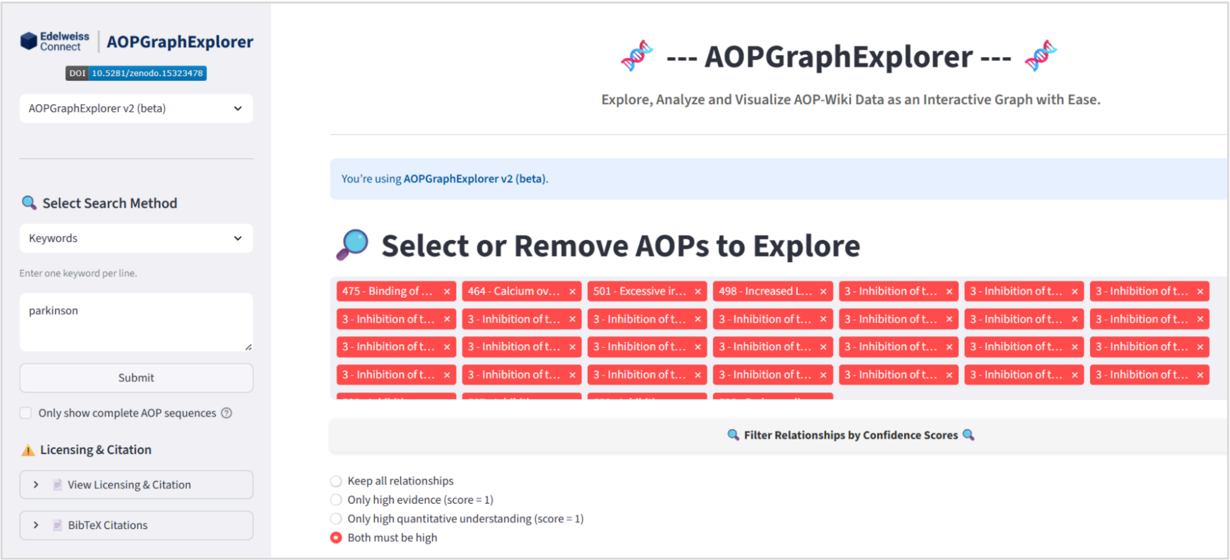


### S3. Network construction and enrichment summary

The integrated network merges AOP events across AOPs by **AOP-Wiki Event ID** and preserves **KER directionality**. Annotation nodes are added as optional overlays and linked to events through typed association edges.

#### Network composition extracted from the export

- **Nodes (85 total)**
  - Molecular Initiating Events (MIE): **12**
  - Key Events (KE): **24**
  - Adverse Outcomes (AO): **3**
  - Processes/GO terms: **27**
  - Phenotypes (HPO/MP): **9**
  - Cell types (CL): **4**
  - Diseases (MeSH): **2**
  - Proteins (PRO): **2**
  - Chemicals (ChEBI): **2**

- **Edges (124 total)**
  - **KER (causal) edges: 52**
  - **Annotation edges: 72** (distributed across several typed relations, including INVOLVES_PROCESS, OBJECT_PARTICIPATES_IN_PROCESS, AFFECTS_GENE_PRODUCT, OCCURS_IN_CELL_TYPE, KE_INVOLVES_CHEMICAL, etc.)

Graph statistics (nodes & edge types)

| AOPs in selection | Total nodes | Total edges | Node types |
|---|---|---|---|
| 9 | 85 | 124 | 9 |

#### Nodes by type

| | Node type | count |
|---|---|---|
| 0 | Process | 27 |
| 1 | KeyEvent | 24 |
| 2 | MolecularInitiatingEvent | 12 |
| 3 | Phenotype | 9 |
| 4 | CellType | 4 |
| 5 | AdverseOutcome | 3 |
| 6 | Protein | 2 |
| 7 | Chemical | 2 |
| 8 | Disease | 2 |

#### Edges by type

| | Edge type | count |
|---|---|---|
| 0 | Key Event Relationship | 52 |
| 1 | OBJECT_PARTICIPATES_IN_PROCESS | 23 |
| 2 | INVOLVES_PROCESS | 18 |
| 3 | KE_INVOLVES_PROCESS | 10 |
| 4 | KE_INVOLVES_OBJECT | 10 |
| 5 | AFFECTS_GENE_PRODUCT | 4 |
| 6 | OCCURS_IN_CELL_TYPE | 4 |
| 7 | KE_INVOLVES_CHEMICAL | 2 |
| 8 | ASSOCIATED_WITH | 1 |

Download node type stats (CSV) Download edge type stats (CSV)

### S4. How to interpret the HTML export (what the reader should do)

The exported HTML provides an interactive network view with built-in interpretation aids:

1. **Node colours** encode node category (AOP events vs annotation layers).
2. **Node size and border thickness** increase with the number of AOPs sharing that node (tooltips explicitly show **"Shared in: X AOPs"** for shared nodes).
3. **KER edges** are directed (arrowheads) and represent causal relationships.
4. **Edge width** increases with the number of AOPs sharing that KER (edge tooltips list contributing **AOP_IDs**).
5. **Click a node** to highlight its local neighbourhood (upstream/downstream context). Tooltips provide rich metadata and outbound links (e.g., AOP-Wiki event/relationship pages; AmiGO links for GO terms).

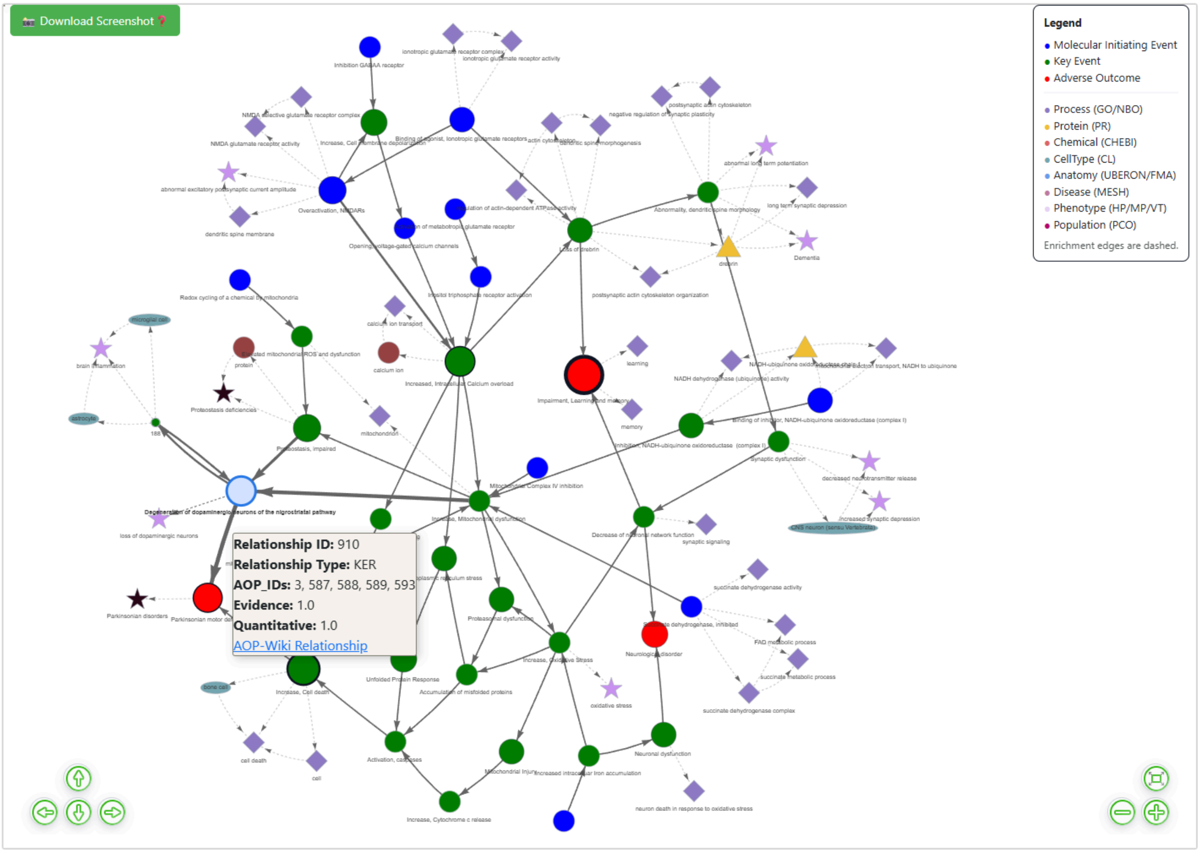
Download Screenshot
Legend
Molecular Initiating Event
Key Event
Adverse Outcome
Process (GO/NBO)
Protein (PR)
Chemical (CHEBI)
CellType (CL)
Anatomy (UBERON/FMA)
Disease (MESH)
Phenotype (HP/MP/VT)
Population (PCO)
Enrichment edges are dashed.
Relationship ID: 910
Relationship Type: KER
AOP_IDs: 3, 587, 588, 589, 593
Evidence: 1.0
Quantitative: 1.0
AOP-Wiki Relationship

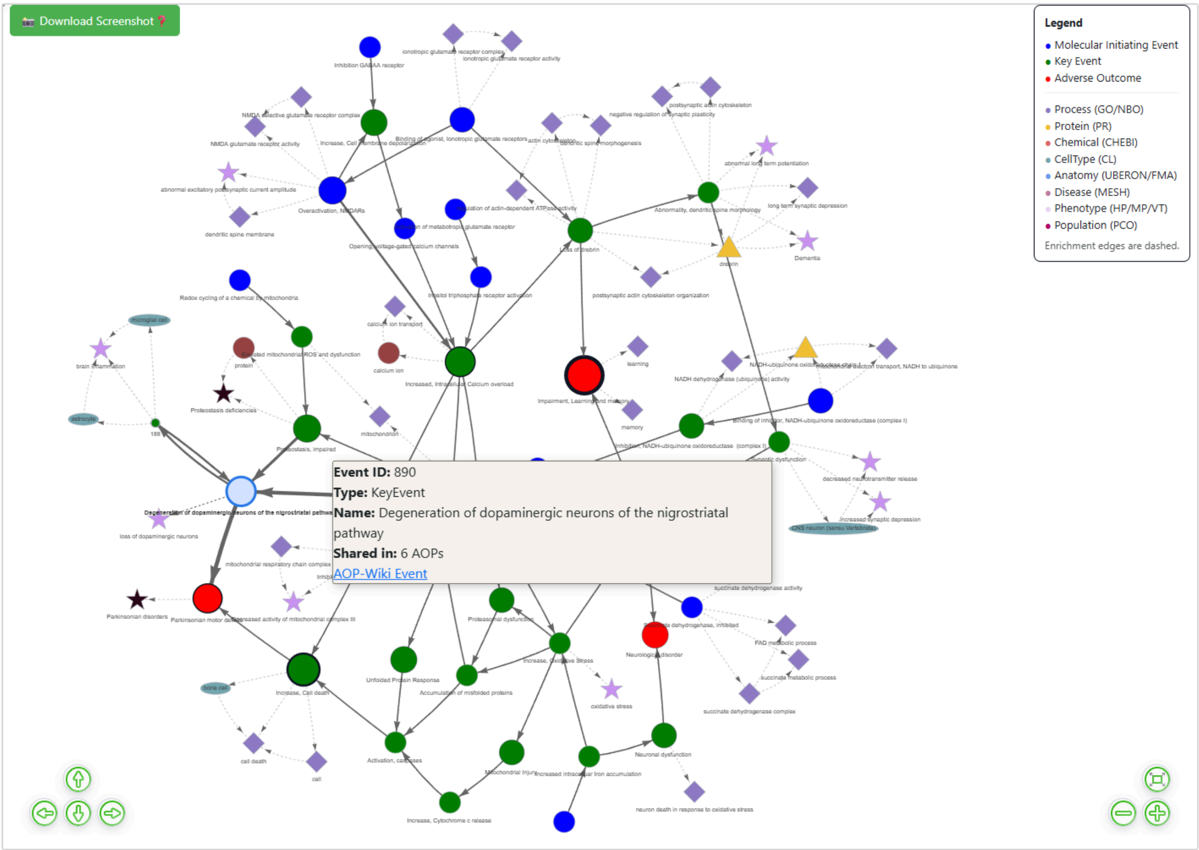
Download Screenshot
Legend
Molecular Initiating Event
Key Event
Adverse Outcome
Process (GO/NBO)
Protein (PR)
Chemical (CHEBI)
CellType (CL)
Anatomy (UBERON/FMA)
Disease (MESH)
Phenotype (HP/MP/VT)
Population (PCO)
Enrichment edges are dashed.
Event ID: 890
Type: KeyEvent
Name: Degeneration of dopaminergic neurons of the nigrostriatal pathway
Shared in: 6 AOPs
AOP-Wiki Event

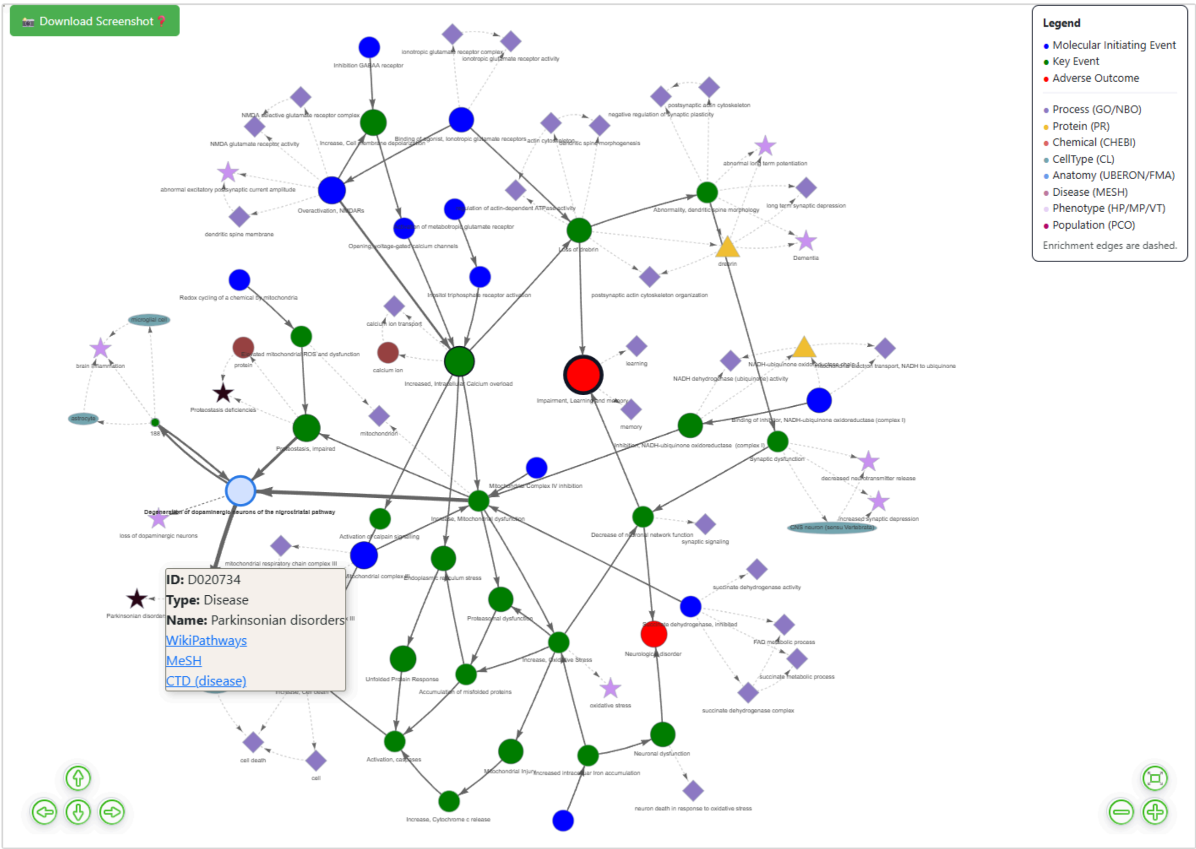
Download Screenshot
Legend
Molecular Initiating Event
Key Event
Adverse Outcome
Process (GO/NBO)
Protein (PR)
Chemical (CHEBI)
CellType (CL)
Anatomy (UBERON/FMA)
Disease (MESH)
Phenotype (HP/MP/VT)
Population (PCO)
Enrichment edges are dashed.
ID: D020734
Type: Disease
Name: Parkinsonian disorders
WikiPathways
MeSH
CTD (disease)

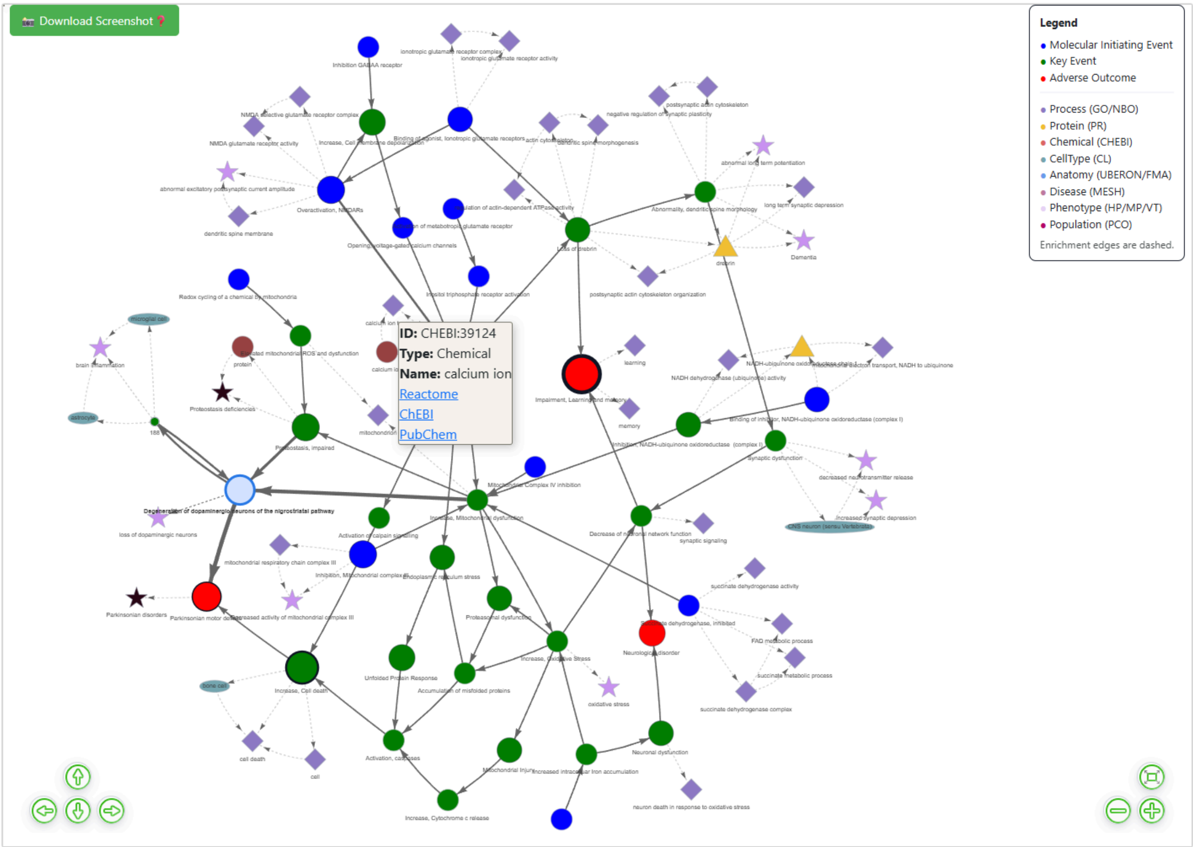
Download Screenshot
Legend
Molecular Initiating Event
Key Event
Adverse Outcome
Process (GO/NBO)
Protein (PR)
Chemical (CHEBI)
CellType (CL)
Anatomy (UBERON/FMA)
Disease (MESH)
Phenotype (HP/MP/VT)
Population (PCO)
Enrichment edges are dashed.
ID: CHEBI:39124
Type: Chemical
Name: calcium ion
Reactome
ChEBI
PubChem

### S5. Evidence-aware filtering demonstration (full network vs high-confidence subnetwork)

KER edges include optional **Evidence** and **Quantitative Understanding** scores in edge tooltips. To demonstrate confidence-based exploration, we compared the **full causal core** against a stricter subnetwork retaining only KERs with **Evidence ≥ 4** and **Quantitative ≥ 4** (where both were reported).

- **Full causal core:** 52 KERs connecting 39 AOP events
- **High-confidence subnetwork:** 24 KERs connecting 19 AOP events

Applying this stricter filter yields a **smaller, more interpretable mechanistic backbone** that emphasizes the most strongly supported relationships and removes many lower-confidence connections that can obscure signal in the integrated view. In the filtered view shown (high-confidence relationships only), the retained causal chain forms a compact, synapse-focused route:

**Binding of agonist to ionotropic glutamate receptors (MIE) → Loss of drebrin (KE) → Impairment in learning and memory (AO)**

Importantly, the **annotation overlays remain available** (dashed enrichment edges), helping contextualize the high-confidence causal path with biologically coherent supporting themes, including **ionotropic glutamate receptor activity/complex**, **dendritic spine morphogenesis**, **postsynaptic actin cytoskeleton organization**, and synaptic plasticity phenotypes such as **abnormal long-term potentiation** and **long-term synaptic depression**, as well as a dementia-related phenotype node. Together, this illustrates how evidence-aware filtering can (i) **prioritize a defensible "core" route** for interpretation and reporting while (ii) still allowing users to understand *why* the retained chain is plausible through surrounding functional and phenotypic context.

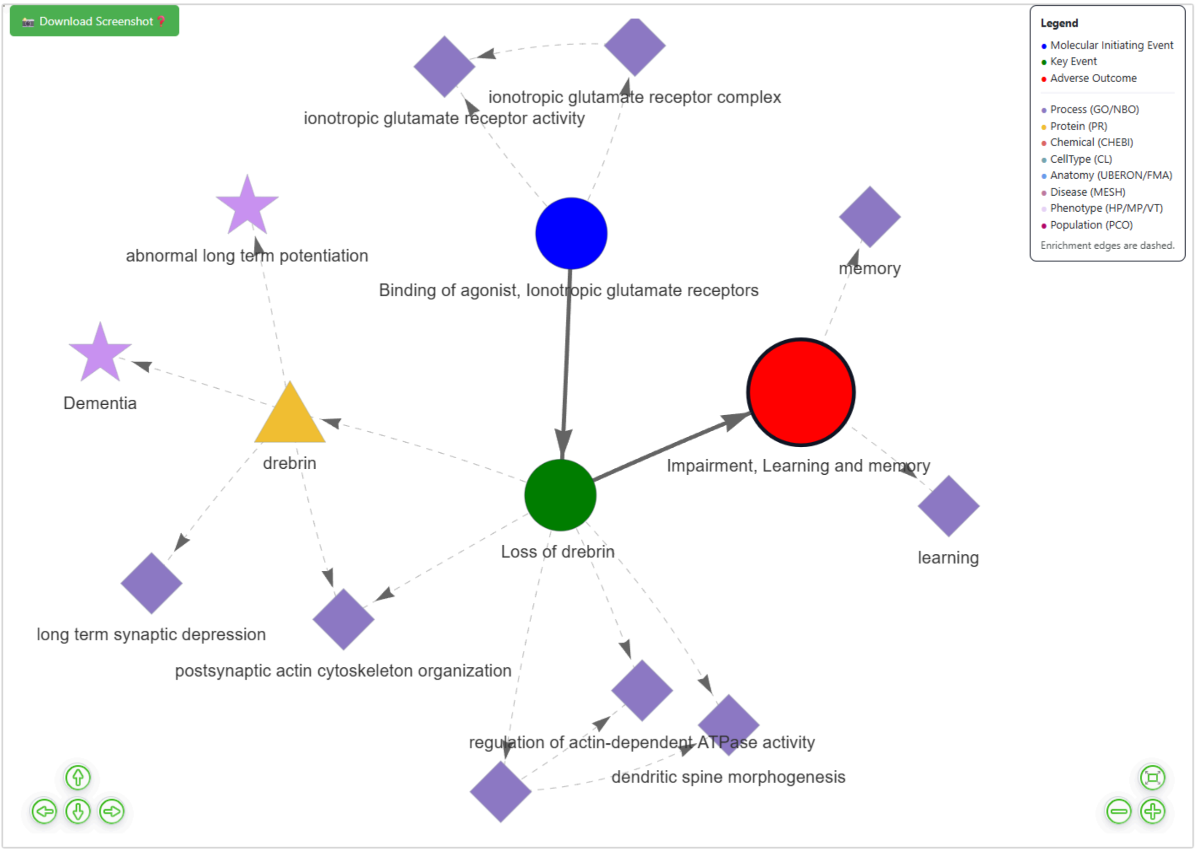


### S6. Mechanistic motifs and representative causal routes to PD-like outcomes

The integrated view supports identification of convergent motifs (multiple upstream perturbations flowing into shared bottlenecks and common adverse outcomes). Below are representative routes to the PD-relevant AO **Parkinsonian motor deficits** (AO ID 896). Relationship IDs and evidence/quant scores are available in the accompanying DOCX tables.

#### Motif 1 — Mitochondrial electron transport inhibition (complex I)

A canonical mitochondrial toxicity route is visible from complex I inhibitor binding through mitochondrial dysfunction to dopaminergic neurodegeneration and motor impairment:

- Binding of inhibitor, NADH-ubiquinone oxidoreductase (complex I)
  → Inhibition of complex I
  → **Increase, mitochondrial dysfunction**
  → **Degeneration of dopaminergic neurons of the nigrostriatal pathway**
  → **Parkinsonian motor deficits**

**Motif 2 — Glutamatergic excitotoxicity / Ca2+ overload (high-confidence core route)**
In the high-confidence subnetwork, a compact route emphasizes excitotoxic calcium loading and downstream cell death:

- Overactivation, NMDARs
  → **Increased, intracellular calcium overload**
  → Activation of calpain signalling
  → **Increase, cell death**
  → **Parkinsonian motor deficits**

This route is useful for prioritizing KEs that represent actionable bottlenecks (e.g., calcium overload, calpain activation, cell death) when focusing on the strongest supported links.

**Motif 3 — Iron accumulation → oxidative stress → proteostasis/cell death**
The network also captures iron-linked oxidative stress and proteostasis stress leading to apoptotic cascades:

- Increased LCN2/iron complex binds SLC22A17 receptor in neuron
  → Increased intracellular iron accumulation
  → **Increase, oxidative stress**
  → Accumulation of misfolded proteins
  → Activation, caspases
  → **Increase, cell death**
  → **Parkinsonian motor deficits**

**Motif 4 — Mitochondrial redox cycling → ROS/dysfunction → proteostasis impairment**
A route consistent with redox cycling stress highlights mitochondrial ROS/dysfunction coupling to proteostasis failure and dopaminergic neurodegeneration:

- Redox cycling of a chemical by mitochondria
  → Elevated mitochondrial ROS and dysfunction
  → **Proteostasis, impaired**
  → **Degeneration of dopaminergic neurons of the nigrostriatal pathway**
  → **Parkinsonian motor deficits**

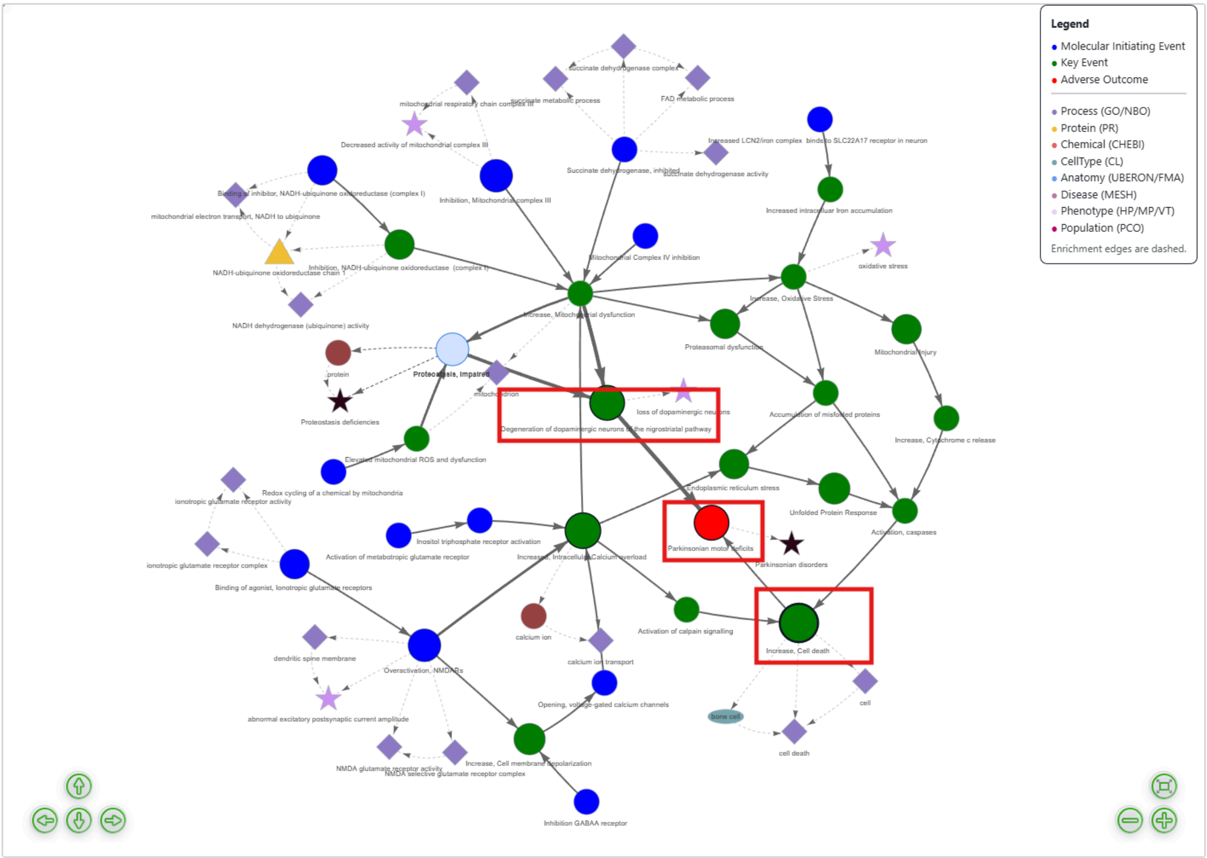


### S7. Shared bottlenecks across AOPs (prioritization signal)

Shared nodes and shared KERs identify convergence points reused across multiple AOPs. In this export, the most shared nodes include:

- **Impairment, learning and memory** (shared across **21** AOPs)
- **Increase, cell death** (shared across **11** AOPs)
- **Increased, intracellular calcium overload** (shared across **7** AOPs)
- **Degeneration of dopaminergic neurons of the nigrostriatal pathway** (shared across **6** AOPs)
- **Parkinsonian motor deficits** (shared across **6** AOPs)

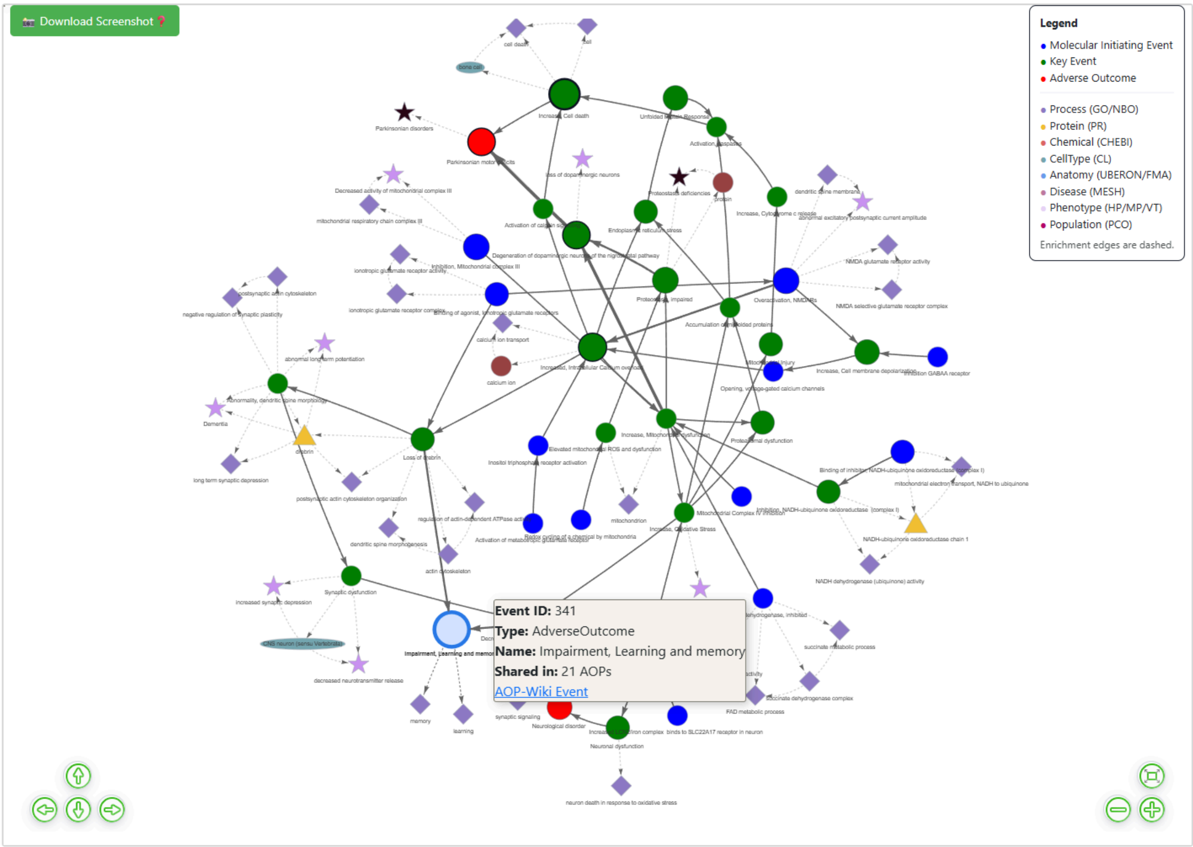


Shared KERs (thicker edges) include the final steps linking dopaminergic neuron degeneration to motor deficits and upstream convergence KERs linking mitochondrial dysfunction to dopaminergic neuron degeneration—supporting the interpretation that these are **cross-AOP bottlenecks** suitable for prioritization.

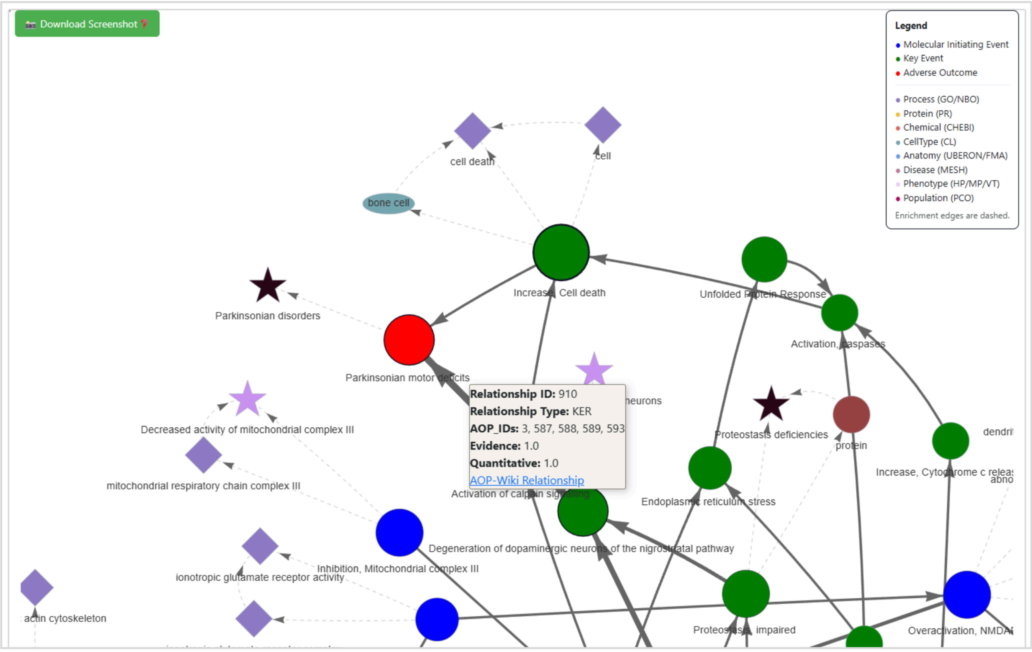

### S8. Annotation-layer examples (how enrichment supports interpretation)

In this export, the enrichment layer includes:

- **Proteins (PRO):** NADH-ubiquinone oxidoreductase chain 1 (PR:000031316), drebrin (PR:000006296), linked via AFFECTS_GENE_PRODUCT edges.
- **Chemicals (ChEBI):** calcium ion (CHEBI:39124) linked to intracellular calcium overload; “protein” (CHEBI:36080) linked to proteostasis impairment.
- **Phenotypes (MP/HPO):** oxidative stress, brain inflammation, loss of dopaminergic neurons, dementia-related terms, synaptic depression/LTP abnormalities.
- **Cell types (CL):** CNS neuron, astrocyte, microglial cell (plus one non-CNS term present in this export).

These overlays help explain *why* specific causal routes are biologically plausible (e.g., calcium ion and calcium transport terms anchoring calcium overload; oxidative stress phenotype anchoring ROS-centric routes).

### S9. Notes and limitations

This analysis reflects the content of the PD-relevant AOP set retrieved from **AOP-Wiki** via keyword-based search and subsequent network assembly in AOPGraphExplorer 2.0. The tool also supports alternative exploration modes, including visualization of **individual AOPs** or aggregation of **all AOPs associated with a selected Key Event**, which may yield different network scope and connectivity depending on the query strategy.

Annotation coverage is inherently **export- and source-dependent**. Enrichment layers (e.g., genes/proteins, chemicals, anatomy/cell types, processes, phenotypes, diseases) rely on the presence of **mappable identifiers and cross-references** in AOP-Wiki entries and the availability of resolvable links in external resources. Therefore, missing annotations in the exported network (e.g., lack of a gene/protein/stressor link for a given event) should be interpreted as **missing or unmapped metadata**, not as evidence that no biological association exists.
